# Whole slide imaging is a high-throughput method to assess *Candida* biofilm formation

**DOI:** 10.1101/2020.03.15.992735

**Authors:** Maximilian W. D. Raas, Thiago P. Silva, Jhamine C. O. Freitas, Lara M. Campos, Rodrigo L. Fabri, Rossana C.N. Melo

## Abstract

New strategies that enable fast and accurate visualization of *Candida* biofilms are necessary to better study their structure and response to antifungals agents. Here, we applied whole slide imaging (WSI) to study biofilm formation of *Candida* species. Three relevant biofilm-forming *Candida* species (*C. albicans* ATCC 10231*, C. glabrata* ATCC 2001, and *C. tropicalis* ATCC 750) were cultivated on glass coverslips both in presence and absence of widely used antifungals. Accumulated biofilms were stained with fluorescent markers and scanned in both bright-field and fluorescence modes using a WSI digital scanner. WSI enabled clear assessment of both size and structural features of *Candida* biofilms. Quantitative analyses readily detected reductions in biofilm-covered surface area upon antifungal exposure. Furthermore, we show that the overall biofilm growth can be adequately assessed across both bright-field and fluorescence modes. At the single-cell level, WSI proved adequate, as morphometric parameters evaluated with WSI did not differ significantly from those obtained with scanning electron microscopy, considered as golden standard at single-cell resolution. Thus, WSI allows for reliable visualization of *Candida* biofilms enabling both large-scale growth assessment and morphometric characterization of single-cell features, making it an important addition to the available microscopic toolset to image and analyze fungal biofilm growth.

## INTRODUCTION

Biofilms are defined as surface-attached, highly resistant, multicellular microbial communities. They are the product of numerous microbes aggregating and some organisms have been observed to wholly merge in this process (Douglas, 2003; Hall-Stoodley, et al., 2004). The main constituents of these remarkably resilient structures when produced by yeast-type fungal species are filamentous hyphae, pseudohyphae, yeast-form cells and a self-produced heterogeneous extracellular matrix (Fanning & Mitchell, 2012; Lagree, et al., 2018).

*Candida* biofilm resilience encompasses several aspects. Firstly, they are highly resistant to antifungal agents. Furthermore, when compared to planktonic phase fungal cells, biofilm infections have increased virulence, higher recurrence rates and are known to readily disperse, forming secondary infection sites (Del Pozo, 2018; Nobile & Johnson, 2015; Ramage, et al., 2012; Uppuluri, et al., 2010). In addition, these pathogenic structures have been shown to suppress immune responses, rendering the host’s intrinsic defences ineffective (Kernien, et al., 2017; Xie, et al., 2012). Taken together, *Candida* biofilms are hard to eradicate and actively promote their survival, procreation and virulence.

As fungal infection disease burden rises alongside increasingly prevailing resistance, research into the fungal biofilm architecture and their response to antifungal agents is becoming more and more pivotal towards successful treatment strategies. A widely used methodology in this research field is scanning electron microscopy (SEM). It provides detailed, ultra-high spatial resolution of the complex three-dimensional (3D) structure that is a biofilm. Other traditional microscopy techniques have been used as well, such as conventional light microscopy and confocal laser scanning microscopy (Lagree, et al., 2018), mainly to study local architecture and molecular markers, respectively (Paulitsch, et al., 2009). Using these techniques, many meaningful insights concerning architecture and mechanisms of resistance have been acquired.

However, traditional microscopy methods are time-consuming and have their limitations when it comes to complete evaluation of large-scale architecture and biofilm response to antifungal agents, as these techniques focus on a very small section of a biofilm from which the whole structure has to be estimated (Azeredo, et al., 2017). Since biofilms are highly complex, multifaceted and heterogeneous (Kean, et al., 2018; Wimpenny, et al., 2000), this limited view will oftentimes lead to interpretative flaws in understanding the complete structure.

In order to obtain more insights into the cohesion of domains inside a biofilm and to investigate whole biofilm architecture normally growing or following exposure to antifungals, we address here the use of a technique new to the field of biofilm research: whole slide imaging (WSI). This approach allows high-resolution, digital visualization of regular glass slides in their entirety with the use of a digital slide scanner thus generating “digital slides” that offer detailed morphological information and possibility for quantification of multiple morphometric parameters (Amaral, et al., 2017; Melo, et al., 2019; Webster & Dunstan, 2014). Other WSI benefits are digital storage, easy share ability, the possibility to make annotations on the slide, automated data extraction, remote accessibility and educational purposes (Saco, et al., 2016).

In the present work, WSI was applied, for the first time, to compare biofilm formation of three species of *Candida (C. albicans, C. glabrata*, and *C. tropicalis*) normally growing in cultures and after treatment with antifungals. Our findings demonstrate that WSI is a powerful tool to assess *Candida* biofilm extent and is able to reveal biofilm morphology even at single-cell level.

## MATERIALS & METHODS

### Fungal strains

The following yeast strains were used in this study: *Candida albicans* ATCC® 10231™*, Candida glabrata* ATCC® 2001™ and *Candida tropicalis* ATCC® 750™. These strains were cultured at 35°C for 24h (*C. albicans* and *C. glabrata*) or 48h (*C. tropicalis*) in Sabouraud dextrose agar (SDA) before all experiments (Campos, et al., 2018).

### Minimal inhibitory concentration (MIC)

The MIC of fluconazole or nystatin was determined by serial microdilution technique according to CLSI-M27 (Rex, 2008). The stock solution of antifungals (fluconazole or nystatin) was diluted in dimethylsulfoxide (DMSO - final concentration of 1%) to obtain concentrations ranging from 100 to 0.78 μg/mL (*C. glabrata* and *C. tropicalis*) or 1,000 to 7.8 μg/mL (*C. albicans*). All tests were performed in a volume of 200 μL and plates were incubated at 35°C for 24h (*C. albicans* and *C. glabrata*) or 48h (*C. tropicalis*). Controls were established without the addition of any antifungal. The MIC values were calculated as the highest dilution showing complete inhibition of tested strain. All analyses were performed in triplicate.

### Biofilm cultures and antimicrobial treatment

Fungal strains were seeded in Sabouraud Dextrose Agar (SDA), incubated (35°C, *C. albicans and C. glabrata*: 24h, C. tropicalis: 48h), and inoculated into a tube with 5 mL of Sabouraud Dextrose Broth (SDB) supplemented with glucose (Aggarwal & Kashyap, 2018; Paiva, et al., 2012). Then, 500 μL of the inoculated broth were added in microtiter plates (24 wells) containing high-adhesion round German glass coverslips (Catalog # 72296-08, EMS, Hatfield, PA, USA), which are used for growing and culturing cells that normally have poor adhesion to regular glass surfaces. For each strain, treatment was performed by adding 500 μL of fluconazole (n=4) or nystatin (n=4) at 1 MIC final concentration per well. Wells without addition of any antifungal served as growth controls (n = 4 for each strain). Sterile controls were done in wells with broth only. *C. albicans* and *C. glabrata* microtiter plates were incubated for 24h at 35°C and *C. tropicalis* plates were incubated for 48h at 35°C.

### Biofilm staining

Two classical fluorescent dyes were used for yeast detection, 4’,6-Diamidine-2’-phenylindole dihydrochloride (DAPI) and Calcofluor White (CW). DAPI is a fluorescent dye that binds selectively to double-stranded DNA and CW is a non-specific fluorophore that binds to cellulose and chitin found in cell walls (Hageage & Harrington, 1984; Kapuscinski, 1995). After incubation, supernatants were removed from wells and coverslips were gently washed with sterile phosphate-buffered saline (0,1 M, pH 7,4 PBS) to remove nonadherent cells. Then, biofilms on coverslips (n= 48, 12 for each strain) were fixed in freshly prepared formaldehyde solution (final concentration 3.7% in 0,1 M, pH 7,4 PBS) for 30 min. Then, the fixative solution was removed and the coverslips were gently washed with sterile distilled water. Biofilm staining was done by covering the coverslips with 10 μL of DAPI (0.01 μg/mL in PBS, Sigma-Aldrich, USA) or CW stain solution (CW M2R [0.001 g/mL] and evans blue [0.0005 g/mL], Sigma-Aldrich, USA) for 5-10 min in the dark. All steps of fixation and staining were done directly in microtiter plate wells. Next, each coverslip was carefully removed from the plates and attached (side opposite to biofilm) on the surface of a regular glass slide using Entellan mounting medium (Merck, Darmstadt, Germany) for subsequent scanning as below.

### Image acquisition

A WSI scanner is a robotic microscope capable of digitizing an entire glass slide, using software to merge or stitch individually captured images into a composite digital image (Webster & Dunstan, 2014). The images generated through this technology are commonly referred to as whole slide images, WSIs, whole slide scans or digital slides (Farahani, et al., 2015). DAPI and CW stained coverslips from the three *Candida* species were digitally scanned in two different modalities (fluorescence and bright-field) using a 3D Scan Pannoramic Histech scanner (3D HistechKft, Budapest, Hungary) connected to a computer (Fujitsu Technology Solutions GmbH, Munich, Germany). High-resolution images (tiff files uncompressed RGB with 300 DPI) obtained from slides scanned at fluorescence (DAPI and CW) and bright-field modes were acquired using Pannoramic Viewer 1.15.2 SP2 RTM software (3D Histechkft Budapest, Hungary) at different digital magnifications (1-100 X).

### Image processing and total biofilm area

Image processing and morphometric analyses of *Candida* biofilms were done using *FIJI* software (National Institutes of Health, Bethesda, MD). First, digital images from CW and DAPI-stained biofilms, scanned at both fluorescent and bright-field modes, were acquired from the entire surface of the glass coverslips at 1-4x digital magnification and converted to binary (black and white, 0-255) images. Next, images were thresholded to highlight just the biofilm areas through pixel saturation. The threshold processing was done in comparison to the original images to ensure the accuracy of all biofilm areas. Then, quantification of the coverslip areas covered by biofilms was performed by measuring the saturated-pixel areas and establishing the proportions of biofilms.

### Scanning electron microscopy

*C. albicans*, *C. glabrata* and *C. tropicalis* biofilms (fluconazole and nystatin-treated and respective controls) on the surface of coverslips were fixed in freshly prepared aldehydes (1% paraformaldehyde and 1.25% glutaraldehyde) in 0.1 M, pH 7.4 phosphate buffer for 1 h at room temperature and processed as before (Lemos, et al., 2018). After post-fixation in 1% osmium tetroxide, the biofilms were washed in phosphate buffer and dehydrated through a graded series of ethanol solutions (30%, 50%, 70%, 90% and twice in 100%) for 15 min at each concentration. Then, samples were critical point dried in carbon dioxide. Subsequently, coverslips were mounted on aluminium holders, sputtered with 5 nm gold and imaged with a double beam scanning electron microscope (Quanta™ 3D FEG, FEI Company, Hillsboro, OR, USA). To study the ultrastructural aspects of the cells, electron micrographs were taken at magnifications of 2,000 – 25,000×.

### Cell morphology analyses

The assessment of *Candida* cellular morphology in biofilms was done by both WSI (cells stained with CW) and SEM. Scanned images (40-100 × digital magnification, RGB or grayscale) from CW stained biofilms and electron micrographs (magnifications of 5,000-25,000×) were analyzed using FIJI software (National Institutes of Health, Bethesda, MD). Cell area and roundness were analyzed for a total of 680 yeast cells as shown in Table 1. The roundness was automatically calculated by 4*area/(π*major_axis^2), with 1.0 indicating a perfect circle and larger values indicating oblong cells.

**Table 1:** Number of *C. albicans, C. glabrata* and *C. tropicalis* cells analyzed by WSI and SEM.

| Experimental Groups | *C. albicans* |  | *C. glabrata* |  | *C. tropicalis* |  |
| --- | --- | --- | --- | --- | --- | --- |
|  | WSI | SEM | WSI | SEM | WSI | SEM |
| Growth control (GC) | 41 | 50 | 46 | 40 | 34 | 33 |
| Fluconazole | 40 | 50 | 36 | 40 | 21 | 35 |
| Nystatin | 34 | 44 | 41 | 28 | 39 | 28 |
| **Total (n)** | **115** | **144** | **123** | **108** | **94** | **96** |

### Statistical Analysis

Data from biofilm areas and cell morphology were compared using Anova followed by Tukey’s comparison test, or Kruskal-Wallis test. The relationship between biofilm areas acquired by different scan modalities (bright-field and fluorescence [DAPI or CW]) was determined using least square regression. All statistics was performed using GraphPad Prism version 8.00 for Windows (GraphPad Software, La Jolla California, www.graphpad.com), considering *P* < 0.05 as a threshold level for significance.

## RESULTS

### MIC analyses

The MIC assay indicates susceptibility or resistance of fungal strains. First, we evaluated the MIC for all fungal strains in response to fluconazole or nystatin as presented in Table 2.

**Table 2:** MIC values for fluconazole and nystatin for the different fungal strains

| *Candida* species | MIC Fluconazole (µg/mL) | MIC Nystatin (µg/mL) |
| --- | --- | --- |
| *C. albicans* | 1,000 | 2.5 |
| *C. glabrata* | 50 | 12.5 |
| *C. tropicalis* | 12.5 | 12.5 |

Strains are considered susceptible to fluconazole if their MIC is <64 μg/mL. Subsequently, resistance is defined at having a MIC of ≥64 μg/mL (Ghannoum & Rice, 1999; Pfaller, et al., 2006). For nystatin, the breakpoint is defined as a MIC of <16 μg/mL (=susceptibility) and ≥16 μg/mL (=resistance) (Kuriyama, et al., 2005). Given this information, *C. albicans* strain can be considered resistant to fluconazole and susceptible to nystatin. The other strains (*C. glabrata* and *C. tropicalis)* were considered susceptible to both fluconazole and nystatin.

### Assessment of biofilms with WSI

Having established the MICs for all strains, cultures of *Candida* species were grown on the surface of round glass coverslips within plates with or without the presence of antifungal agents (Fig. 1). After the incubation period, coverslips were stained with two different fluorophores [DAPI (n=24) and CW (n=24)], affixed on glass slides and scanned in a digital scanner with three different WSI strategies: bright-field (BF) mode (all slides, n=48), fluorescence mode after DAPI staining (n=24) and fluorescence mode after CW staining (n=24) (Figs 1A-D). Thus, a total of 96 high-resolution virtual images covering the total biofilm area were acquired for subsequent viewing and morphometric analyses using software (Figs 1E, F). Each virtual image represents an entire glass coverslip/slide (Fig. 1). Comparisons were drawn between BF and fluorescence modalities to explore their accuracy and reliability.

**Figure 1 -.**
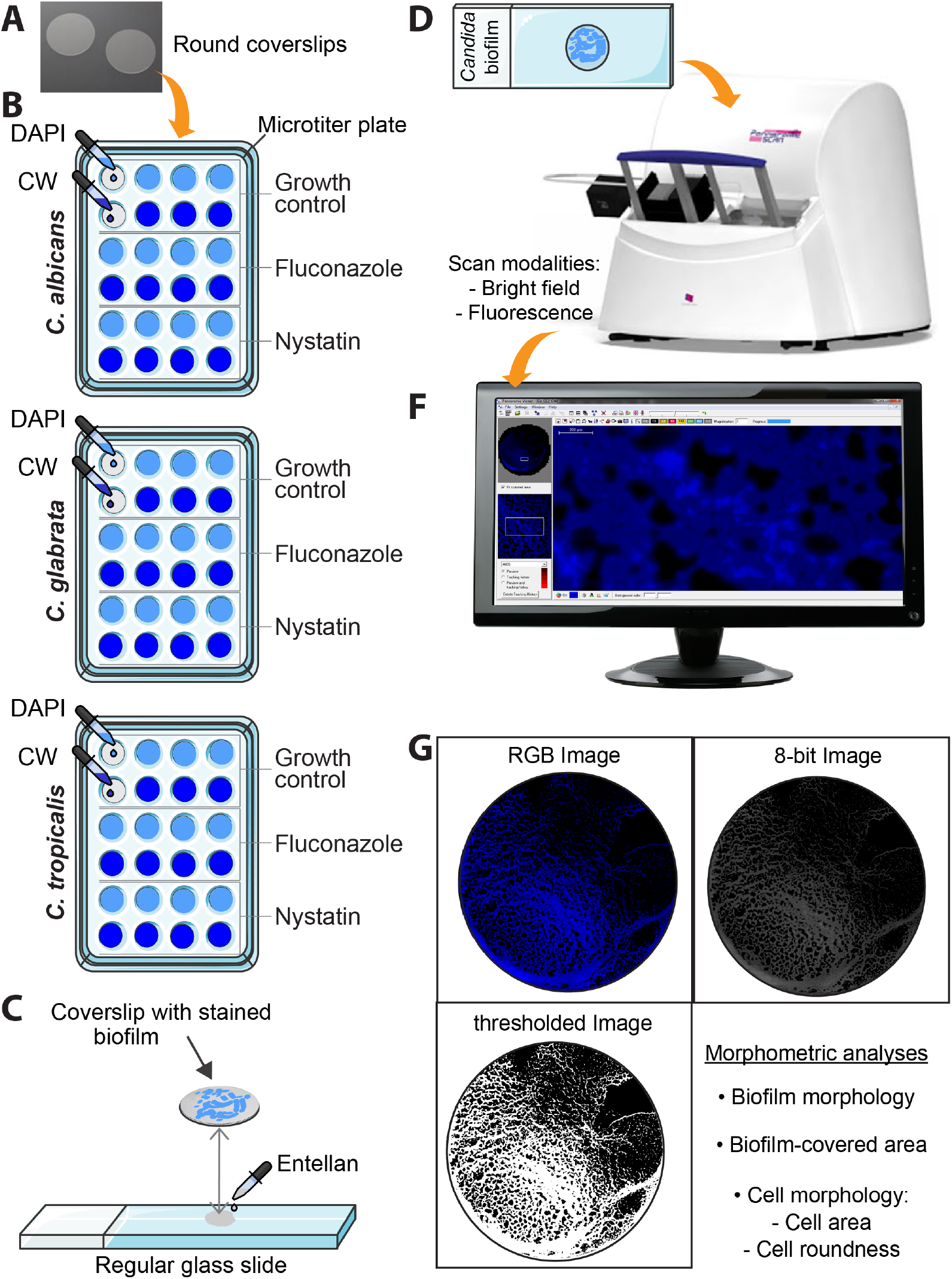
Assessment of biofilms with WSI. (A) Round glass coverslips were used for growing biofilms in cultures. (B) Cultures of *C. albicans, C. glabrata* and *C. tropicalis* were cultivated in microtiter plates with or without the presence of antifungals agents and developed biofilms on the surface of coverslips were stained with two different fluorophores [DAPI and Calcofluor-white (CW); n=4/group/each staining]. (C) Stained coverslips were affixed on regular glass slides using Entellan mounting medium. (D) Glass slides with coverslips were scanned in a digital scanner (3D HistechKft, Budapest, Hungary) in two different modalities (bright-field and fluorescence). (E) High-*resolution images were acquired using Pannoramic Viewer 1.15.2 SP2 RTM software (3D Histechkft* Budapest, Hungary). (F) virtual images (RGB) were converted to 8-bit (black and white) and thresholded for morphometric analyses using FIJI software.

### WSI enables reliable quantification of biofilm development

The biofilm surface area is a reliable measure of the fungal ability to attach to a surface and proliferate (Rex, 2008). Here, we used coverslips made with German glass, which provide high adhesion of cells in cultures and are indicated for optimal imaging with fluorescent probes, to successfully grow Candida biofilms and then applied WSI. In growth control samples, we clearly notice the formation of consistent biofilms attached on the surface of coverslips (Figs 2 A-C). Considering all scan strategies together (bright field and fluorescence), our quantitative analyses showed that GC biofilms covered more than 80% of the coverslips surface (Figs 2 D-F). Thus, WSI showed to be a very useful tool to examine large-scale biofilms of *Candida* species, both in normal growth and in the presence of antifungal agents, as shown below for each species.

**Figure 2 -.**
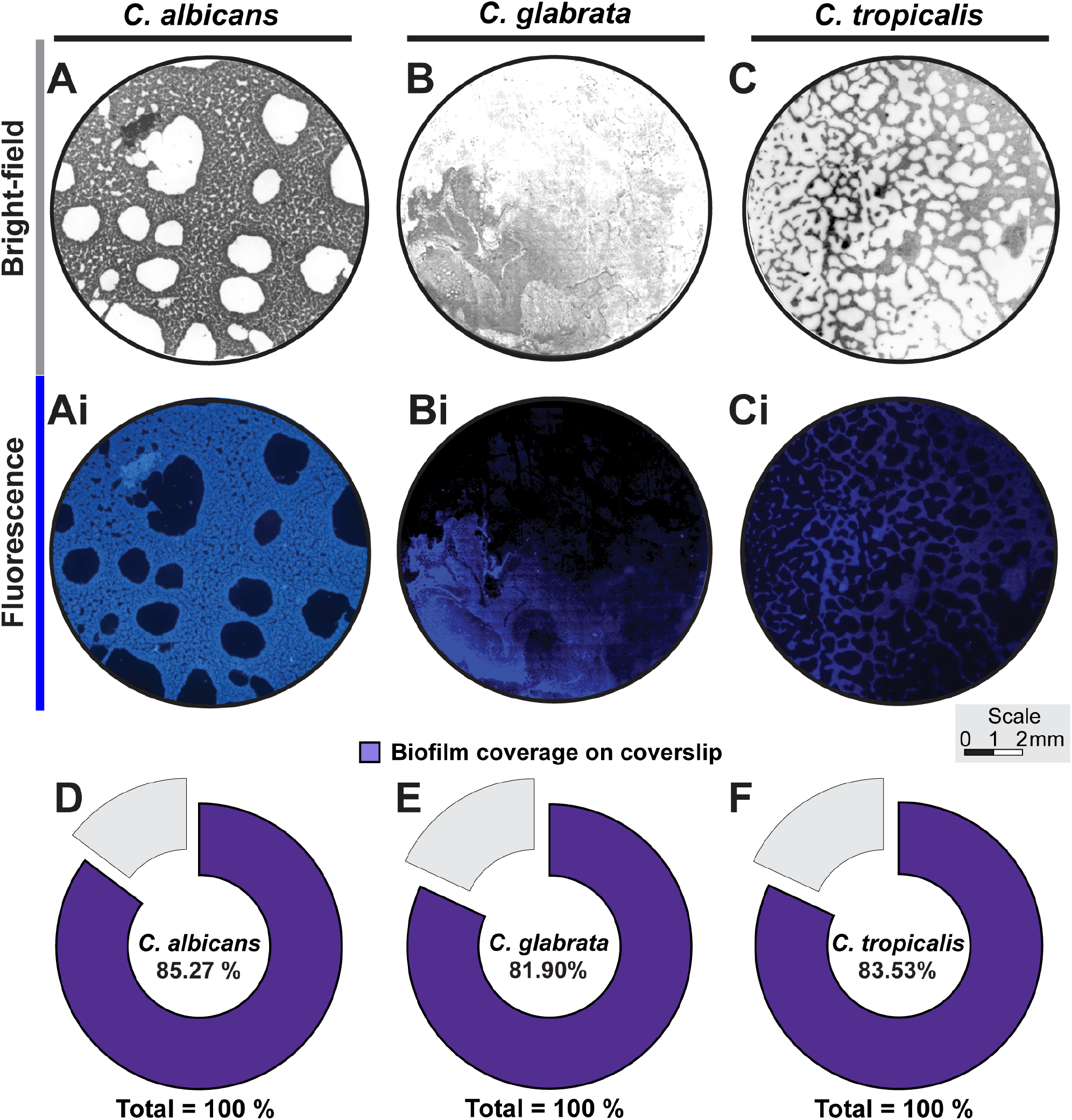
Evaluation of *Candida* biofilms growth with WSI. Representative virtual images of *C. albicans* (A and Ai), *C. glabrata* (B and Bi) and *C. tropicalis* (C and Ci) biofilms growing at the surface of coverslips in the control group acquired under bright-field and fluorescence microscopy. In (D-F), the proportion of biofilm-covered area is shown considering all scan modes. A total of 96 high-resolution virtual images covering the total biofilm area were acquired for subsequent viewing and morphometric analyses using FIJI software. Each virtual image represents an entire glass coverslip/slide. Fluorescent images show biofilms stained with DAPI.

#### C. albicans

Compared to the control group (Figs 2A, Ai), fluconazole exposure did not affect biofilm formation and adhesion of the fluconazole-resistant *C. albicans* across the coverslips (Fig. 3 A). On the other hand, nystatin-exposed *C. albicans* cells were not able to establish any meaningful biofilm on the coverslip (Figs 3 A, D). WSI quantitative analyses in all WSI modalities showed a significant decrease of biofilm-covered area in the presence of antifungals (. 3F; *P*<0.0001 for all). When the extent of biofilms acquired with different WSI modalities were compared, there were no significant differences among all three strategies (Fig. 3F; *P*>0.999 for all).

**Figure 3 -.**
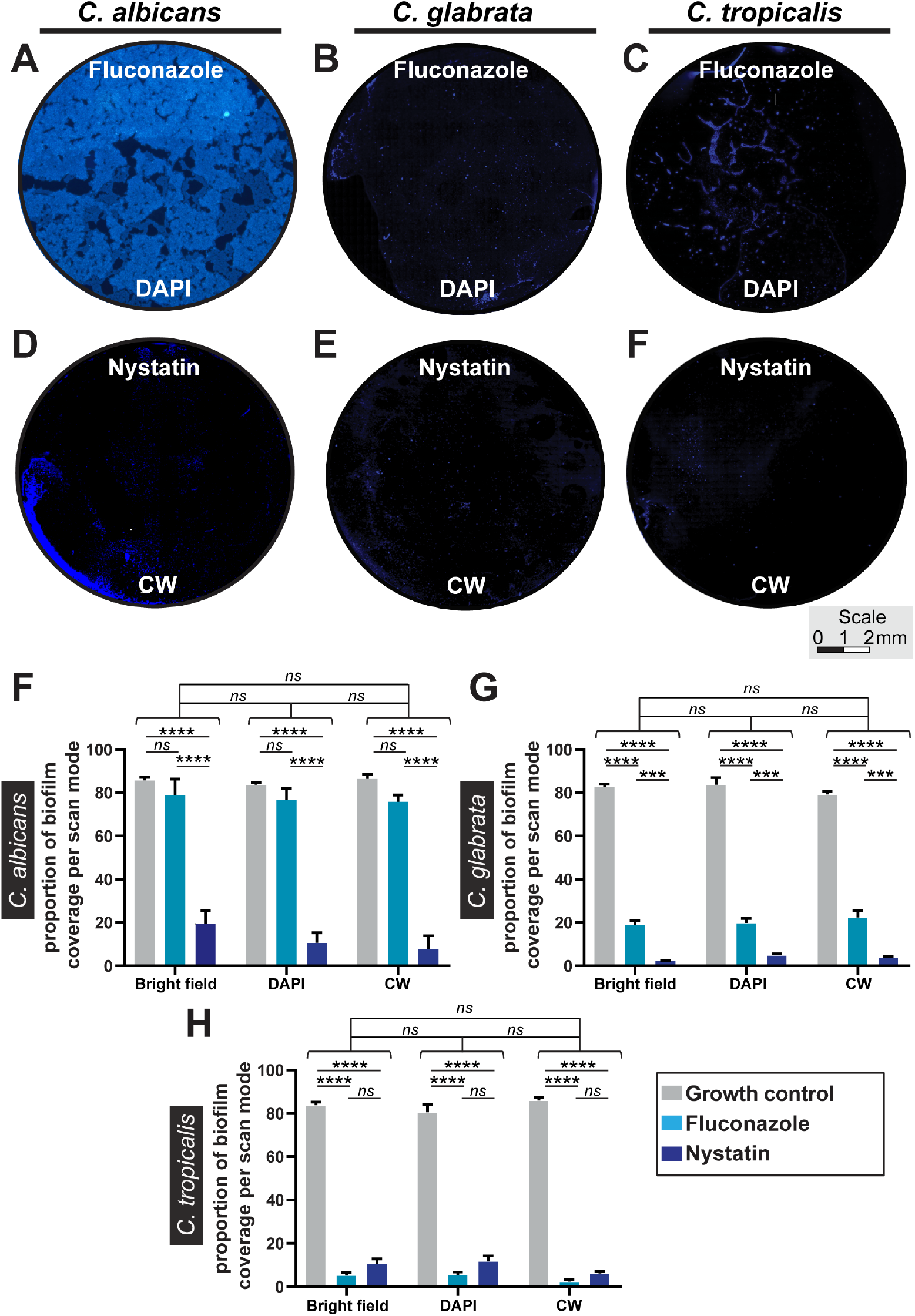
Evaluation of *Candida biofilms* formation after fluconazole or nystatin treatments. (A-I) Virtual images of biofilms growing at the surface of coverslips in the control group (A-C) and after antifungal treatments (D-I) acquired with different scans strategies. In (J), the biofilm-covered areas were quantitated for each group considering all scan modes. (K) The proportion of biofilm coverage is shown per each scan mode. Significant (asterisks, *P*<0.0001) and non-significant (NS) differences are indicated when groups were compared.

#### C. glabrata

This *Candida* strain was susceptible to both antifungals. This was clearly demonstrated by the extent of formed biofilms. While in the control group (Figs 2A, Ai), biofilms occupied most of the coverslip surface, treated samples displayed very low levels of biofilm coverage (Figs 3B, E). Measurements of the biofilm-covered areas taken in all scanning modalities confirmed the drastic decrease in the amount of biofilm formed following exposure to antifungals compared to the control group (Fig. 3G; *P*<0.0001). As shown for *C. albicans,* individual WSI modes did not affect visualization of biofilm areas (Fig. 3G; *P*>0.999 for all). In each WSI modality, a significant decrease in the proportion of biofilm coverage was observed after fluconazole or nystatin treatments (BF, DAPI, CW: *P*<0.0001 for each).

#### C. tropicalis

This *Candida* strain showed no resistance during the MIC experiments. Accordingly, compared to the control group (Figs 2A, Ai), large decreases in the biofilm extent were observed by *C. tropicalis* in the presence of fluconazole or nystatin (Fig. 3C, F; *P*<0.0001). Quantitative analyses in all WSI modes confirmed a significant reduction in the proportion of biofilm-covered areas after both treatments in comparison to the control group (Fig. 3H; *P*<0.0001). Each individual WSI mode also detected significant reduction of the biofilm areas in response to antifungals (BF, DAPI, CW: *P*<0.0001).

Of note, application of an additional statistical method (regression analysis) for all *Candida* species and treatments demonstrated that the total surface area of biofilms measured under different WSI modalities were correlated thus demonstrating that the biofilm areas can be reliably detected by BF and fluorescence modes (Fig. S1).

### WSI is a useful tool to assess biofilm structure

By measuring the biofilm areas visualized with WSI, we noticed that additional information on structural aspects of *Candida* biofilms could be obtained from the high-resolution images when seen in high magnification. Thus, WSI revealed the organization of the biofilms for the different *Candida* species (Fig. 4). *C. albicans* generally clustered in one large and highly dense biofilm with channel-like structures flowing through them (Figs 4A, B). This aspect was not seen for *C. glabrata* (Figs 4C, D) *and C. tropicalis* (Fis. 4E, F) biofilms, which appeared as a less dense structure with typical meshwork aspect. While *C. glabrata* biofilm is characterized by thin filamentous stretches of relatively small fungal cells surrounding an empty, almost round opening (Figs 4C, D), *C. tropicalis* (Figs 4E, F) forms a meshwork with thicker septa with a slightly lesser cell density. These septa also include circular openings of their own creating an intricate pattern presented.

**Figure 4 -.**
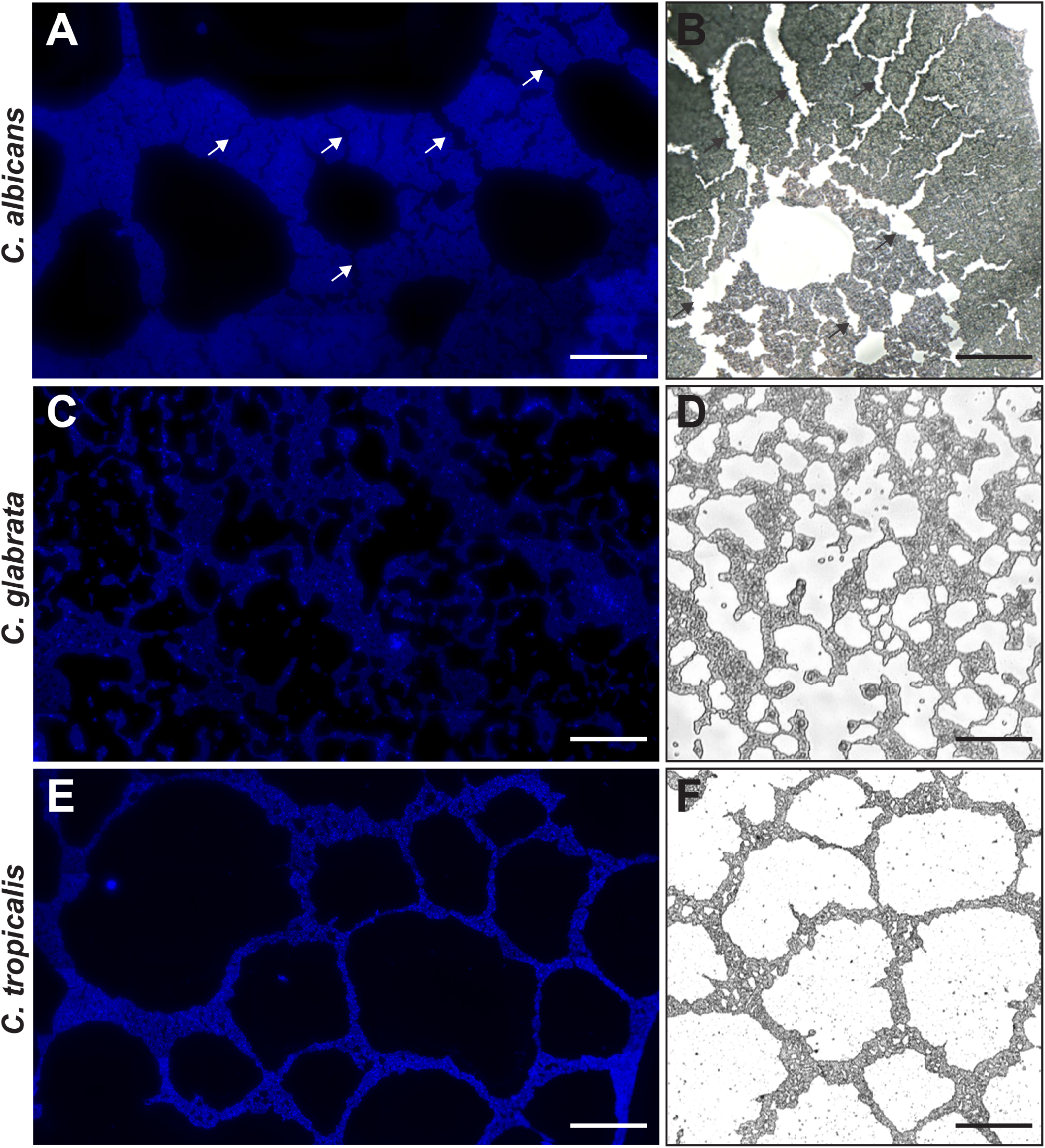
Architecture of *Candida* biofilms normally growing in cultures. (A, B) *C. albicans* form a highly dense biofilm with channel-like structures (arrows) flowing through the biofilm extent. (C, D) *C. glabrata* biofilm is characterized by a meshwork shape with thin filamentous stretches of relatively small fungal cells surrounding an empty, almost round opening. (E, F) *C. tropicalis* biofilm shows a honeycomb-like chambers. WSI images were acquired under bright-field (A, C, F) or fluorescence/DAPI staining (B, D, F) modes. Scale bar, 200 μm.

WSI also showed the presence of filamentous cells (hyphal forms) in biofilms of all *Candida* species (Figs 5A-C), indicative of vegetative growth through a filamentation process, which is considered a virulence factor (Berman, 2006). Hyphal forms are frequently present in the later latter stages of fungal biofilms maturation and can represent pseudohyphae or true hyphae in *Candida* biofilms (Roilides & Ramage, 2017). Division and constriction septa between cells in the filament were clearly observed when the samples were stained with CW (Fig. 5A).

**Figure 5 -.**
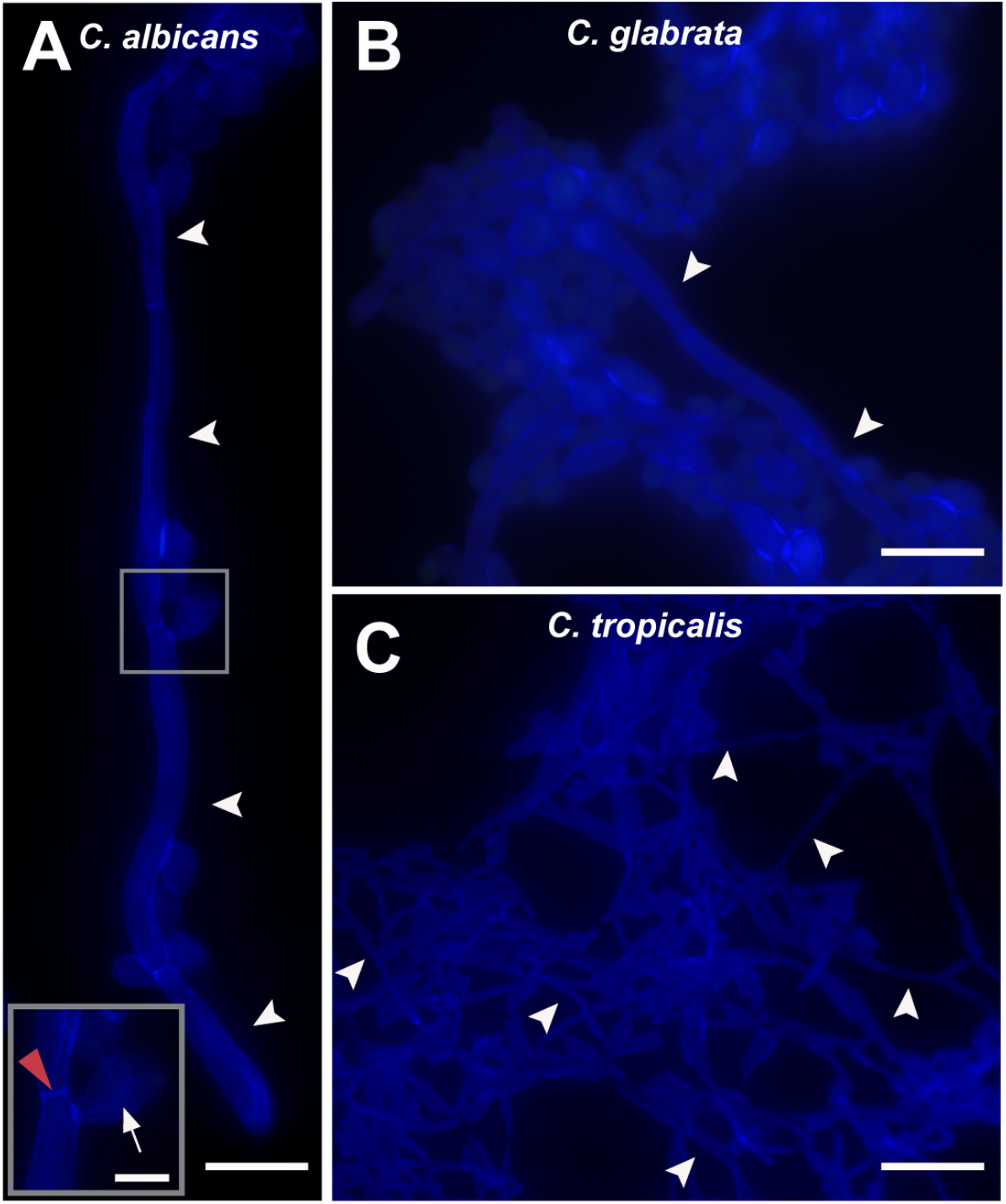
Hyphal forms with typical morphology (arrowheads) are seen in *C. albicans* (A), *C. glabrata* (B) and *C. tropicalis* (C) biofilms. In (A), the presence of constriction septa separating filamentous cells is indicated by a red arrowhead in higher magnification in the inset. Budding cells (arrow) are also noted. Scale bar, 15 μm.

### WSI reveals biofilm morphology at single-cell level

WSI revealed not only the species-specific organization of the biofilms (Fig. 4), but also morphological aspects (cell shape) of individual yeast cells. High-resolution WSI analyses of biofilms stained with CW, which enables visualization of the cell surface (cell wall), enabled the observation of round and elongated yeasts cells in both RGB (Figs 6A-C) and grayscale (Figs 6D-F).

**Figure 6 -.**
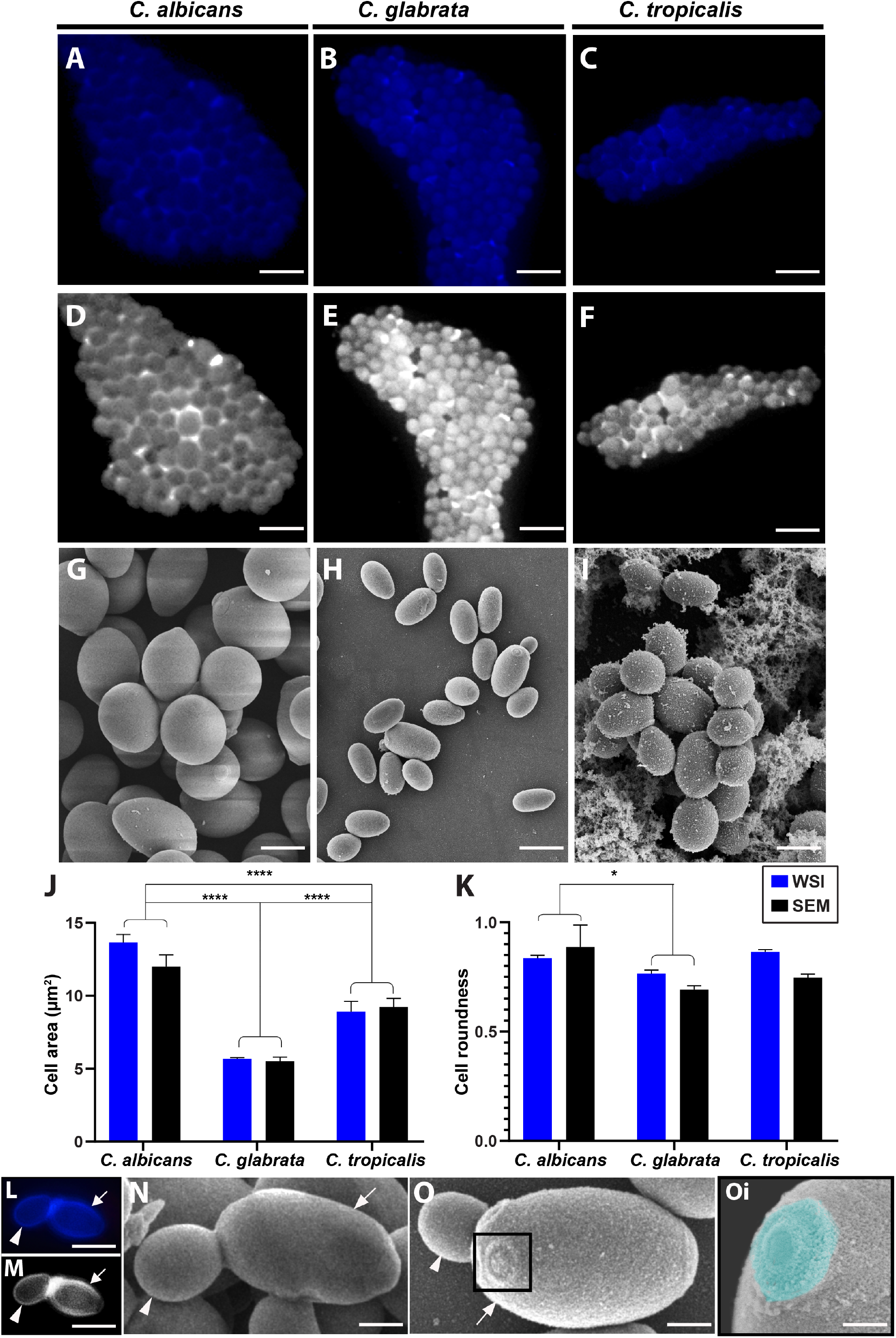
Single-cell morphology of *Candida* biofilms normally growing in cultures. (A-C) High-resolution WSI images of *C. albicans, C. glabrata* and *C. tropicalis*, respectively, observed at bright-field mode. In (D-F), the same fields are observed by WSI in fluorescence mode. In (G-I), images from scanning electron microscopy (SEM) shows the same shapes observed by WSI. Note in (I) that part of the fungal extracellular matrix is observed in the background. (J) Cell area and (K) cell roundness were quantitated in images acquired with WSI and SEM, with no difference between these two tools. In (L-O) observe by WSI (L and M) or SEM (N-O) the presence of budding cells (arrowheads) originated from mother yeast cells (arrows). Note in (Oi) a bud scar (highlighted in cyan) on the surface of a yeast cell. A total of 332 WSI images and 348 electron micrographs were analysed. Asterisk indicates significant difference (*P* = 0.01). Scale bar; 6 μm (A-F), 3 μm (G-I), 4 μm (L-M), 800 nm (N, O), 400 nm (Oi). Asterisk indicates significant difference (J: *P* <0.0001; K: *P* = 0.01).

To evaluate the accuracy of biofilm-composed yeast cell shapes, we next prepared additional samples of *Candida* species growing in normal conditions to SEM, a technique considered to be the “gold standard” to study cell morphology in biofilms (Azeredo, et al., 2017). SEM (Figs 6G-I) showed in 3D the same morphological aspects depicted by WSI images (Fig. 6A-F). We then applied quantitative analyses to evaluate cell area and cell roundness in the images acquired by both WSI and SEM. These two parameters are considered relevant to study yeast cell shapes (Bougen-Zhukov, et al., 2017; Lee, et al., 2011; Liu, et al., 2011; Wang & Lin, 2012). In all cases, the data for cell morphometric assessment with WSI did not differ from the data acquired with SEM (*P* >0.90 for all), indicating that the resolution of WSI is accurate at this level and for this purpose.

Remarkably, morphometric analyses revealed that the area of individual yeast cells in the biofilm significantly varies depending on the *Candida* species and that this variation can be detected by both WSI and SEM (Figs 6J, *P* < 0.0001). Cell roundness analyses, done with both WSI and SEM images, revealed significant differences when *C. albicans* was compared to *C. glabrata* (Fig. 7K, *P* = 0.01). Additionally, cell budding was clearly identified by both WSI (Figs 6L, M) and SEM (Figs 6N, O) in all three species in process of vegetative growth. Both WSI and SEM revealed that while mother cells exhibited a more elongated shape, budding cells had a pronounced round shape and smaller size (Figs 6L-N). “Bud Scars” on the surface of some yeast cells were seen in detail by SEM (Figs 6O and 6Oi). Evidence of budding was more frequently observed in *C. glabrata*, which could explain their reduced area and roundness in comparison with *C. albicans* and *C. tropicalis* (Figs 6J and 6K).

## DISCUSSION

Most fungal pathogens exist as adhered cells to surfaces within a well-organized biofilm ecosystem and not as free-floating organisms. However, the architecture of fungal biofilms is not fully understood. In the present study, we explore the use of WSI as an approach for large-scale imaging of *Candida* biofilms. One of the advantages of WSI is that it produces images with high resolution and, at the same time, with a wide field of observation. Here, we show that *Candida* species were able to attach and grow as biofilms on coverslips, as expected (Malm, et al., 2010; Silva-Dias, et al., 2015) and that WSI application facilitated comprehensive visualization of *Candida* biofilms both in natural growth and in presence of antifungal agents. Moreover, WSI combined with computational methods enabled fast quantification of the adhesion extent of *Candida* biofilms. Thus, WSI showed to be a very useful tool to examine large-scale biofilms of *Candida* species, since in growth control samples biofilms covered most surfaces of the coverslips.

Our data show that inhibitory concentrations of antifungal agents applied to planktonic fungi can effectively inhibit the formation of biofilms. However, MIC concentration applied to a *C. albicans* resistant-strain did not led to significant decreases in the amount of biofilm formed by this strain within a 24h period as expected. Both biofilm formation and biofilm reduction after treatment with antifungals were properly detected by WSI. Our findings demonstrated no variation in the reduction of area covered by biofilms after treatments when WSI modalities were compared. Thus, WSI can be a valuable tool to accurately test biofilm responses to antifungals regardless if the images are acquired under bright-field or fluorescence (DAPI and CW) WSI modes, indicating reliability of WSI settings.

Remarkably, not only the extent but also structural aspects of biofilms were captured by WSI. This approach revealed, for example, that *C. albicans* form complex and highly organized channels which appear as a ramifying system of channels inside its biofilm that, to our knowledge, have been scarcely studied. As these channels are important size-determining aspects of a biofilm due to their facilitation of nutrient-flow (Ramage, et al., 2001), WSI can be particularly useful to address future studies focused on these structures. Moreover, our approach with WSI enabled detection of hyphal forms (Fig. 5) in all three *Candida* species even in the absence of a specific induction of this process. Filamentous growth, although not strictly essential for biofilm formation, provides protection and adhesion sites for the budding yeast cells and contributes to the overall robustness of the biofilm (de Barros, et al., 2020; Nobile & Johnson, 2015; Roilides & Ramage, 2017). Other structural components revealed by WSI such as division and constriction septa between cells of the filament forms are associated with pseudohyphal formation in *Candida* species (Berman, 2006).

Since WSI delivers high-resolution images, it also offers the possibility to perform high amount of analyses at the single-cell level, thus extracting more biologically significant cellular information. This has been the case of WSI applications to pathological analyses of biopsies in different fields. For example, in the field of liver pathology and biology, WSI has been extensively used for different purposes, such as to understand cellular diversity of hepatic progenitors, scoring of inflammatory cells and identification of specific hepatocarcinoma cell alterations [reviewed in (Campos, et al., 2019)].

Here, we expand the application of WSI to understand morphological aspects of *Candida* species within a biofilm. Our data showed that WSI is a reliable tool for evaluation of the cell area and cell roundness yeast cells as addressed by fluorescence WSI modality. Cellular area and roundness are important parameters related to environmental conditions and can even reflect specific yeast molecular features, such as cell cycle phase, genetic expression (phenotypic characteristics) or virulence (Bougen-Zhukov, et al., 2017; Lee, et al., 2011; Liu, et al., 2011; Wang & Lin, 2012). Our quantitative analyses revealed that these parameters can vary among *Candida* species (Fig. 6) and therefore WSI can be beneficial to understand cell behavior within a biofilm. Yet, the morphometric data acquired from WSI scans showed no significant differences from data extracted from SEM, a technique widely used for biofilm studies and accepted as particularly accurate at single-cell resolution (Azeredo, et al., 2017). Moreover, by observing *Candida* biofilms by WSI, we detected cell budding, which is a common division process for both haploid (*C. glabrata*) and diploid (*C. albicans* and *C. tropicalis*) yeast cells of *Candida* species (Branduardi, et al., 2012), which was confirmed by SEM.

WSI was also found to be a practical approach for imaging *Candida* biofilms. The digitalization is fully automated, and wide-field images can be stored. Complete slide scanning means that every slide can be entirely revisited at any point allowing more detailed assessment as the analyses are not confined to limited fields-of-view captured with a regular light microscope fitted with a camera. Easy accessibility, shareability and storage allowed us to explore and quantify data from the scans using adequate software. The possibility to make annotations on slides was also very convenient, since this procedure highlights specific sections of the scan thus facilitating discussion, including remote access. On the other hand, by our experience, WSI does have some limitations for biofilm studies, especially when using unfixed biofilms stained with fluorochromes, which does not allow adequate focus. The scanning process of fluorescent samples also takes more time compared to bright-field modality, but this issue has been addressed with new generation of scanners.

Finally, our group is the first to apply WSI to understand fungal biofilms produced by different species of Candida with perspective of applications to other fungal biofilms. We recently showed that WSI is helpful to test the potential antifungal action of a phytocompound against a multidrug-resistant *C. tropicalis* strain (Lemos, et al., 2020). WSI can be also useful to study bacterial biofilms. This technique enabled to examine biofilms produced by *Salmonella enterica* strains after treatment with plant extracts with potential anti-bacterial activity (Campos, et al., 2019) and to investigate antibiotic resistance at single-bacterium level (Song, et al., 2019).

In conclusion, WSI is an emerging technique that proved to be a powerful tool in delivering an accurate, high-resolution and full-length view of *Candida* biofilms and therefore can be a worthy and efficient addition to the available microscopic toolset to image and analyse fungal biofilms.

## Author’s contributions

RM provided the study design, study mentorship and critical editing of the manuscript. MR, TS and RF contributed to the study design. MR and TS performed experiments and acquired and analyzed the data. JF and LC performed cultures. All authors contributed in part to writing and editing the manuscript and approved the final version.

## Acknowledgements

The authors would like to thank Andreia Alvim for helpful discussions as well as suggestions made during the scanning electron microscopy analyses. We thank the UFMG Microscopy Center (Centro de Microscopia da UFMG, Belo Horizonte, MG, Brazil) for technical support in sample preparation and image acquisition for scanning electron microscopy.

## Supporting information

**Figure S1-.**
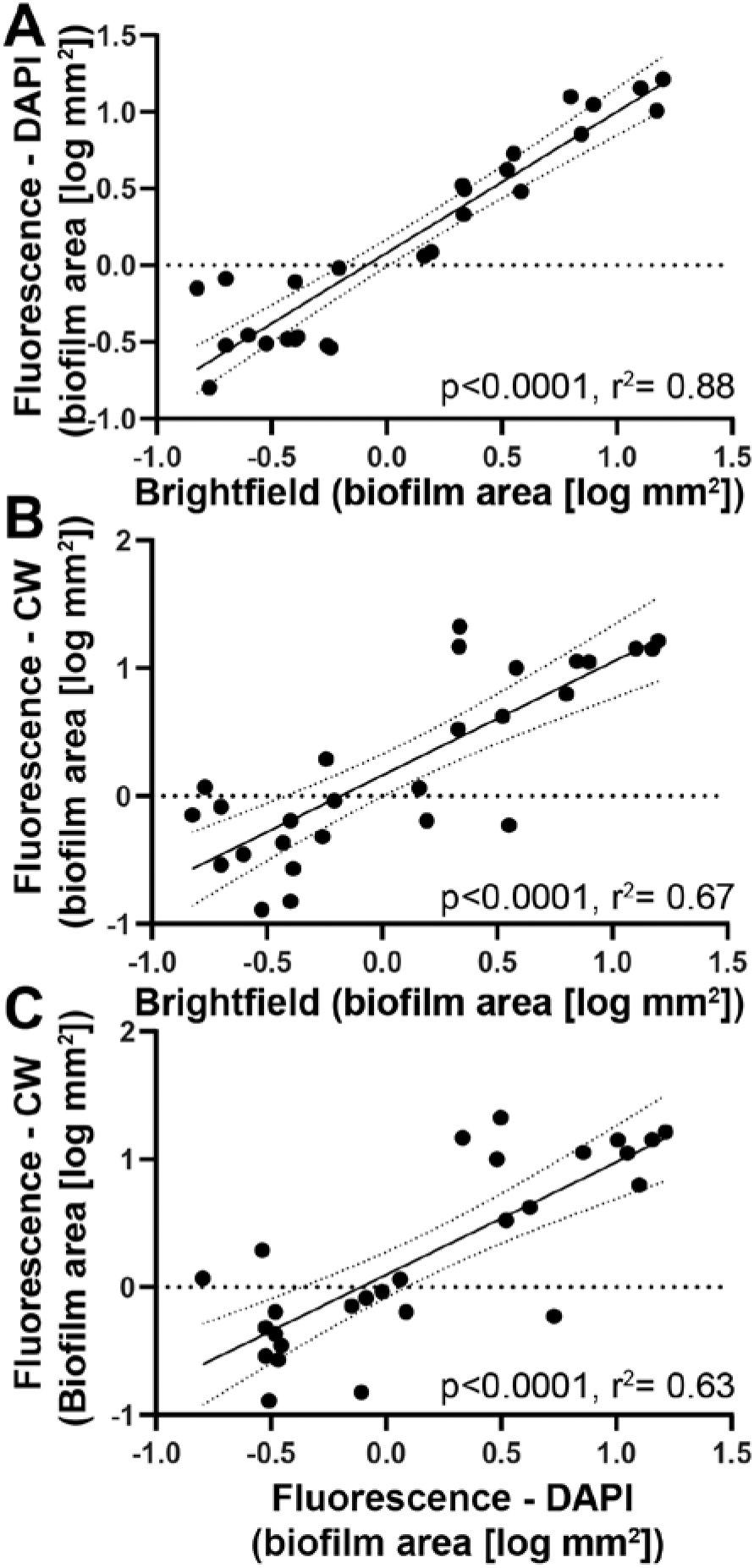
Relationship between biofilm area acquired by different scan modalities and staining methods. DAPI and Calcofluor White (CW)-stained coverslips from three biofilm-forming *Candida* species (*C. albicans* ATCC 10231, *C. glabrata* ATCC 2001, and *C. tropicalis* ATCC 750) were digitally scanned in two different modalities (fluorescence and bright-field). Regression analyses between covered-biofilm area were statistically significant (*P*<0.0001) when methods were compared: (A) Brigh-tfield *versus* Fluorescence - DAPI, (B) Bright-field *versus* Fluorescence - CW and (C) Fluorescence - DAPI *versus* Fluorescence - CW.

## References

Aggarwal, P. & Kashyap, B. (2018). Biofilm production by clinically isolated Candida: Comparative analysis based on specimen, methodology, and various Candida species. Indian J Med Special 9(2), 69–72.

Amaral, K.B., Silva, T.P., Dias, F.F., Malta, K.K., Rosa, F.M., Costa-Neto, S.F., Gentile, R. & Melo, R.C.N. (2017). Histological assessment of granulomas in natural and experimental *Schistosoma mansoni* infections using whole slide imaging. PLoS One 12(9), e0184696.

Azeredo, J., Azevedo, N.F., Briandet, R., Cerca, N., Coenye, T., Costa, A.R., Desvaux, M., Di Bonaventura, G., Hebraud, M., Jaglic, Z., Kacaniova, M., Knochel, S., Lourenco, A., Mergulhao, F., Meyer, R.L., Nychas, G., Simoes, M., Tresse, O. & Sternberg, C. (2017). Critical review on biofilm methods. Crit Rev Microbiol 43(3), 313–351.

Berman, J. (2006). Morphogenesis and cell cycle progression in *Candida albicans*. Curr Opin Microbiol 9(6), 595–601.

Bougen-Zhukov, N., Loh, S.Y., Lee, H.K. & Loo, L.H. (2017). Large-scale image-based screening and profiling of cellular phenotypes. Cytometry A 91(2), 115–125.

Branduardi, P., Dujon, B., Feldmann, H., Gaillardin, C. & Porro, D. (2012). V: Yeast growth and the yeast cell cycle. Molecular and Cell Biology. Feldmann H.(ur.). Weinheim, Wiley-Blackwell, 175–204.

Campos, L.M., Lemos, A.S., Silva, T.P., Oliveira, L.G., Nascimento, A.L.R., Carvalho, J.J., de Moraes, A.C., Rocha, V.N., Aguiar, J.A., Scio, E., Apolônio, A.C., Melo, R.C.N. & Fabri, R. (2019). *Mitracarpus frigidus* is active against *Salmonella enterica* species including the biofilm form. Ind Crops Prod 141, 111793.

Campos, L.M., Melo, L., Lemos, A.S.O., Guedes, M.C.M.R., Silva, T.P., Fig.Ueiredo, G.F., Reis Junior, F.B., Rocha, V.N., Melo, R.C.N., Araújo, M.G.F., Apolonio, A.C.M., Scio, E. & Fabre, A. (2018). *Mitracarpus frigidus:* A promising antifungal in the treatment of vulvovaginal candidiasis. Ind Crops Prod 12, 731–739.

de Barros, P.P., Rossoni, R.D., de Souza, C.M., Scorzoni, L., Fenley, J. & Junqueira, J. (2020). Candida Biofilms: An update on Developmental Mechanisms and Therapeutic Challenges. Mycopathologia 185, 415–420..

Del Pozo, J.L. (2018). Biofilm-related disease. Expert Rev Anti Infect Ther 16(1), 51–65.

Douglas, L.J. (2003). Candida biofilms and their role in infection. Trends Microbiol 11(1), 30–36.

Fanning, S. & Mitchell, A.P. (2012). Fungal biofilms. PLoS Pathog 8(4), e1002585.

Farahani, N., Parwani, A.V. & Pantanowitz, L. (2015). Whole slide imaging in pathology: advantages, limitations, and emerging perspectives. Pathol Lab Med Int 7, 23–33.

Ghannoum, M.A. & Rice, L.B. (1999). Antifungal agents: mode of action, mechanisms of resistance, and correlation of these mechanisms with bacterial resistance. Clin Microbiol Rev 12(4), 501–517.

Hageage, G.J. & Harrington, B.J. (1984). Use of calcofluor white in clinical mycology. Lab Med 15(2), 109–112.

Hall-Stoodley, L., Costerton, J.W. & Stoodley, P. (2004). Bacterial biofilms: from the natural environment to infectious diseases. Nat Rev Microbiol 2(2), 95–108.

Kapuscinski, J. (1995). DAPI: a DNA-specific fluorescent probe. Biotech Histochem 70(5), 220–233.

Kean, R., Delaney, C., Rajendran, R., Sherry, L., Metcalfe, R., Thomas, R., McLean, W., Williams, C. & Ramage, G. (2018). Gaining Insights from Candida Biofilm Heterogeneity: One Size Does Not Fit All. J Fungi (Basel) 4(1).

Kernien, J.F., Snarr, B.D., Sheppard, D.C. & Nett, J.E. (2017). The Interface between Fungal Biofilms and Innate Immunity. Front Immunol 8, 1968.

Kuriyama, T., Williams, D.W., Bagg, J., Coulter, W.A., Ready, D. & Lewis, M.A. (2005). In vitro susceptibility of oral Candida to seven antifungal agents. Oral Microbiol Immunol 20(6), 349–353.

Lagree, K., Desai, J.V., Finkel, J.S. & Lanni, F. (2018). Microscopy of fungal biofilms. Curr Opin Microbiol 43, 100–107.

Lee, H.N., Magwene, P.M. & Brem, R.B. (2011). Natural variation in CDC28 underlies morphological phenotypes in an environmental yeast isolate. Genetics 188(3), 723–730.

Lemos, A.S., Florêncio, J.R., Pinto, N.C., Campos, L.M., Silva, T.P., Grazul, R.M., Pinto, P.F., Tavares, G.D., Scio, E. & Apolônio, A.C.M. (2020). Antifungal activity of the natural coumarin scopoletin against planktonic cells and biofilms from a multidrug-resistant *Candida tropicalis* strain. Front Microbiol 11, 1525.

Lemos, A.S.O., Campos, L.M., Melo, L., Guedes, M., Oliveira, L.G., Silva, T.P., Melo, R.C.N., Rocha, V.N., Aguiar, J.A.K., Apolonio, A.C.M., Scio, E. & Fabri, R.L. (2018). Antibacterial and Antibiofilm Activities of Psychorubrin, a Pyranonaphthoquinone Isolated From *Mitracarpus frigidus (Rubiaceae)*. Front Microbiol 9, 724.

Liu, Y., Aubrey, W., Martin, K., Sparkes, A., Lu, C. & King, R.D. (2011). The analysis of yeast cell morphology features in exponential and stationary Phase. J Biol Syst 19(04), 561–575.

Malm, A., Chudzik, B., Piersiak, T. & Gawron, A. (2010). Glass surface as potential in vitro substratum for Candida famata biofilm. Annals of Agricultural and Environmental Medicine 17(1), 115–118.

Melo, R.C.N., Raas, M.W.D., Palazzi, C., Neves, V.H., Malta, K.K. & Silva, T.P. (2019). Whole Slide Imaging and Its Applications to Histopathological Studies of Liver Disorders. Front Med (Lausanne) 6, 310.

Nobile, C.J. & Johnson, A.D. (2015). *Candida albicans* Biofilms and Human Disease. Annu Rev Microbiol 69, 71–92.

Paiva, L.C., Vidigal, P.G., Donatti, L., Svidzinski, T.I. & Consolaro, M.E. (2012). Assessment of in vitro biofilm formation by Candida species isolates from vulvovaginal candidiasis and ultrastructural characteristics. Micron 43(2-3), 497–502.

Paulitsch, A.H., Willinger, B., Zsalatz, B., Stabentheiner, E., Marth, E. & Buzina, W. (2009). In-vivo Candida biofilms in scanning electron microscopy. Med Mycol 47(7), 690–696.

Pfaller, M.A., Diekema, D.J. & Sheehan, D.J. (2006). Interpretive breakpoints for fluconazole and Candida revisited: a blueprint for the future of antifungal susceptibility testing. Clin Microbiol Rev 19(2), 435–447.

Ramage, G., Rajendran, R., Sherry, L. & Williams, C. (2012). Fungal biofilm resistance. Int J Microbiol 2012, 528521.

Ramage, G., Vandewalle, K., Wickes, B.L. & Lopez-Ribot, J.L. (2001). Characteristics of biofilm formation by *Candida albicans*. Rev Iberoam Micol 18(4), 163–170.

Rex, J.H. (2008). Reference method for broth dilution antifungal susceptibility testing of filamentous fungi: approved standard. Clinical and Laboratory Standards Institute.

Roilides, E. & Ramage, G. Current perspectives in fungal biofilms. In MYCOSES, pp. 13–14. WILEY 111 RIVER ST, HOBOKEN 07030-5774, NJ USA.

Saco, A., Bombi, J.A., Garcia, A., Ramirez, J. & Ordi, J. (2016). Current Status of Whole-Slide Imaging in Education. Pathobiology, 83(2-3), 79–88.

Silva-Dias, A., Miranda, I.M., Branco, J., Cobrado, L., Monteiro-Soares, M., Pina-Vaz, C. & Rodrigues, A.G. (2015). In vitro antifungal activity and in vivo antibiofilm activity of cerium nitrate against Candida species. J Antimicrob Chemother 70(4), 1083–1093.

Song, D., Liu, H., Ji, H. & Lei, Y. (2019). Whole Slide Imaging for High-Throughput Sensing Antibiotic Resistance at Single-Bacterium Level and Its Application to Rapid Antibiotic Susceptibility Testing. Molecules 24(13), 2441.

Uppuluri, P., Chaturvedi, A.K., Srinivasan, A., Banerjee, M., Ramasubramaniam, A.K., Kohler, J.R., Kadosh, D. & Lopez-Ribot, J.L. (2010). Dispersion as an important step in the *Candida albicans* biofilm developmental cycle. PLoS Pathog 6(3), e1000828.

Wang, L. & Lin, X. (2012). Morphogenesis in fungal pathogenicity: shape, size, and surface. PLoS Pathog 8(12), e1003027.

Webster, J.D. & Dunstan, R.W. (2014). Whole-slide imaging and automated image analysis: considerations and opportunities in the practice of pathology. Vet Pathol 51(1), 211–223.

Wimpenny, J., Manz, W. & Szewzyk, U. (2000). Heterogeneity in biofilms. FEMS Microbiol Rev 24(5), 661–671.

Xie, Z., Thompson, A., Sobue, T., Kashleva, H., Xu, H., Vasilakos, J. & Dongari-Bagtzoglou, A. (2012). Candida albicans biofilms do not trigger reactive oxygen species and evade neutrophil killing. J Infect Dis 206(12), 1936–1945.

